# Fission yeast Pak1 phosphorylates anillin-like Mid1 for spatial control of cytokinesis

**DOI:** 10.1101/722900

**Authors:** Joseph O. Magliozzi, Jack Sears, Lauren Cressey, Marielle Brady, Hannah E. Opalko, Arminja N. Kettenbach, James B. Moseley

**Affiliations:** Department of Biochemistry and Cell Biology, The Geisel School of Medicine at Dartmouth, Hanover, NH 03755, USA; Norris Cotton Cancer Center, The Geisel School of Medicine at Dartmouth, Lebanon, NH 03756, USA

## Abstract

Protein kinases direct polarized growth by regulating the cytoskeleton in time and space, and could play similar roles in cell division. We found that the Cdc42-activated polarity kinase Pak1 colocalizes with the assembling contractile actomyosin ring (CAR) and remains at the division site during septation. Mutations in *pak1* led to defects in CAR assembly and genetic interactions with cytokinesis mutants. Through a phosphoproteomic screen, we identified novel Pak1 substrates that function in polarized growth and cytokinesis. For cytokinesis, we found that Pak1 regulates the localization of its substrates Mid1 and Cdc15 to the CAR. Mechanistically, Pak1 phosphorylates the Mid1 N-terminus to promote its association with cortical nodes that act as CAR precursors. Defects in Pak1-Mid1 signaling lead to misplaced and defective division planes, but these phenotypes can be rescued by synthetic tethering of Mid1 to cortical nodes. Our work defines a new signaling mechanism driven by a cell polarity kinase that promotes CAR assembly in the correct time and place.

**Summary:** Magliozzi et al. show that fission yeast cell polarity kinase Pak1 regulates cytokinesis. Through a phosphoproteomic screen and subsequent mutant analysis, their work uncovers direct targets and mechanisms for Pak1 activity during cell division.

## Introduction

Cell polarization and cytokinesis share a requirement for the assembly of actin filaments into higher-order structures that are physically linked to the cell cortex. For example, polarized actin cables, actomyosin stress fibers, and dendritic actin networks are central to cell polarity in yeast and metazoan cells (Tojkander et al., 2012; Chiou et al., 2017; Omotade et al., 2017). The architecture of these actin structures is controlled by cell polarity signaling proteins, which relay spatial landmarks to the cytoskeleton. Yeast and metazoan cells dramatically reorganize the actomyosin cytoskeleton into a contractile actomyosin ring (CAR) at cytokinesis. Past work has shown that cortical cell polarity proteins indirectly regulate cytokinesis by positioning the mitotic spindle, which provides spatial signals CAR assembly (Bringmann and Hyman, 2005; Rappaport et al., 2009; Siller and Doe, 2009; Kotak and Gönczy, 2013). However, there are hints of more direct links between cell polarity proteins and the CAR. For example, cell polarity factors position the CAR in fly neuroblasts that are induced into cytokinesis in the absence of a mitotic spindle (Cabernard et al., 2010), and cortical cell polarity factors contribute to the robustness of cytokinesis in *C. elegans* embryos (Davies et al., 2016; Jordan et al., 2016). In this study, we investigated the possibility of a direct role for polarity signaling factors during cytokinesis of fission yeast cells.

Fission yeast has served as a long-standing model system for studies on both cell polarity and cytokinesis. Genetic screens identified conserved polarity factors essential for the cylindrical shape of fission yeast cells (Hayles and Nurse, 2001). An early screen isolated a class of round (or “orb”) mutants that were enriched for protein kinases including Pak1, also called Orb2 or Shk1 (Snell and Nurse, 1994; Verde et al., 1995; Marcus et al., 1995). PAK kinases are auto-inhibited until activation by binding to the GTP-bound form of the GTPase Cdc42, a global regulator of cell polarity (Bokoch, 2003). Active PAK kinases then organize the cytoskeleton by phosphorylating downstream substrates. In fission yeast, Pak1 phosphorylates several proteins *in vitro* (Yang et al., 1999; Endo et al., 2003; Kim et al., 2003; Yang et al., 2003; Loo and Balasubramanian, 2008), but we do not have a full understanding of Pak1 targets in cells and their functional connections to cell polarity and/or cytokinesis.

At mitotic entry, the actin machinery shifts to the fission yeast cell middle to form the CAR through a tightly regulated process. Many proteins that participate in these events have been identified through decades of genetic screens and subsequent characterization (Chang et al., 1996; Wu et al., 2006; Pollard and Wu, 2010). Fission yeast cytokinesis occurs in four steps. First, a subset of cytokinesis proteins is recruited to 50-75 cortical spots called nodes, which are positioned in the cell middle (Chang et al., 1996; Sohrmann et al., 1996; Wu et al., 2006). The anillin-related protein Mid1 serves as the primary anchor to position cytokinetic nodes in the cell middle (Sohrmann et al., 1996; Paoletti and Chang, 2000; Celton-Morizur et al., 2006; Padte et al., 2006; Almonacid et al., 2009; Almonacid et al., 2011). The second step is CAR assembly, when nodes coalesce into an intact ring through actin-myosin based interactions (Wu et al., 2006; Vavylonis et al., 2008). The third step is CAR maturation, when additional cytokinesis proteins are recruited to the intact ring (Wu et al., 2003; Wu et al., 2006; Pollard and Wu, 2010). The fourth step is ring constriction, when the CAR constricts through the combined forces of actin-myosin interactions and cell wall deposition (Liu et al., 1999; Proctor et al., 2012; Ramos et al., 2019).

Recent studies identified a direct role for the cell polarity kinases Pom1 and Kin1 in fission yeast cytokinesis. Kin1 and Pom1 phosphorylate multiple cytokinesis proteins including shared substrates (Kettenbach et al., 2015; Lee et al., 2018; Bhattacharjee et al., 2020). Both *in vitro* and in cells, Kin1 and Pom1 phosphorylated largely non-overlapping residues on these shared substrates (Lee et al., 2018). Simultaneous inhibition of both kinases led to cytokinesis defects, including “unspooling” of the CAR during constriction (Lee et al., 2018). These results led us to hypothesize that additional cell polarity kinases might directly regulate cytokinesis proteins to promote cell division. Here, we screened for cell polarity kinases that localize to the site of cytokinesis and found a new role for Pak1 in CAR assembly. Through large-scale phosphoproteomics, we identified novel Pak1 substrates including Mid1. Phosphorylation by Pak1 promotes Mid1 association with cortical node precursors for proper cytokinesis. Our work reveals a new function for this conserved kinase in cytokinesis and supports the growing role of cell polarity signaling in cytokinesis.

## Results and Discussion

### Cdc42-Pak1 localizes to the assembling CAR

To identify cell polarity kinases that might function in cytokinesis and cell division, we monitored the localization of seven mEGFP-tagged kinases during cell division. Each strain also expressed the myosin light chain Rlc1-mRuby2 to mark the CAR and Sad1-mCherry to mark the spindle-pole body (SPB), which allowed us to categorize kinase localization during defined stages of cell division (Fig. 1A). Six of the kinases were recruited to the cell middle late in the division process, after CAR assembly and during constriction. In contrast, Pak1 localized to the cell middle shortly after SPB splitting, when CAR node precursors were still coalescing into a ring. Upon closer examination by Airyscan confocal microscopy, Pak1 colocalized with CAR marker Rlc1 during ring assembly, but we also observed Pak1 signal at the division site that did not overlap with Rlc1. Interestingly, Pak1 formed a strand that linked two loose ends of the CAR in the final stages of assembly (Fig. 1B). No such strands were observed in the presence of Latrunculin B (Fig. S1A), but we note Pak1 colocalization with F-actin marker Lifeact was only partial during CAR assembly (Fig. S1B). These results suggest that Pak1 might function during CAR assembly.

**Figure 1:**
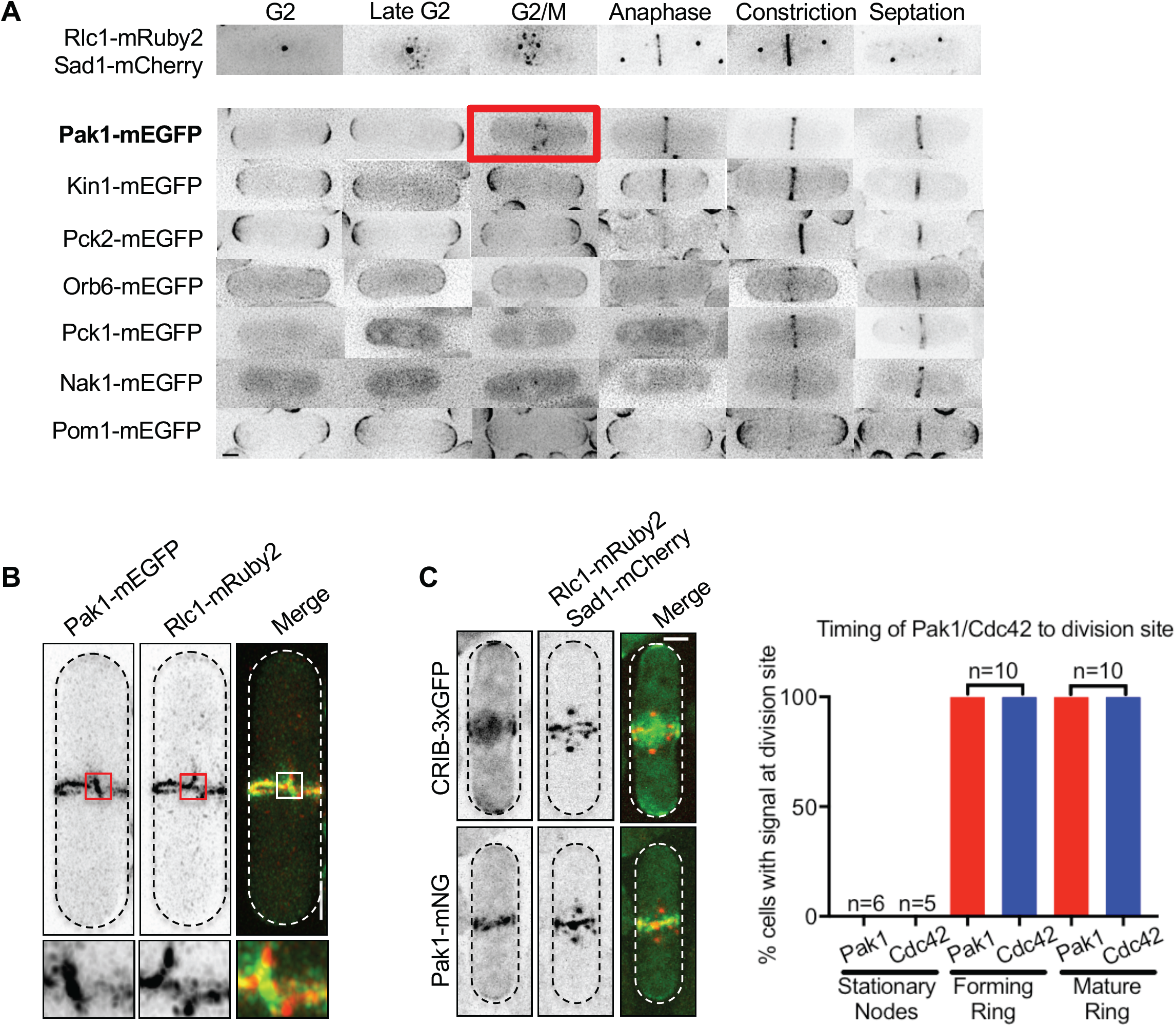
Cell polarity kinase Pak1 and activated Cdc42 localize to the cell division site. **(A)** Localization of the indicated protein kinases during defined stages of cell division. Red box highlights Pak1-mEGFP localization during CAR assembly. **(B)** Colocalization of Pak1-mEGFP and Rlc1-mRuby2 by Airyscan confocal microscopy. Insets are enlarged views of the boxed regions. Images are maximum intensity projections. **(C)** Left: Localization of Pak1 and Cdc42 biosensor at the assembling CAR. Right: quantification of CAR localization. Scale bars, 2 μm.

Pak1 is a member of the PAK (p21-activated protein kinase) family and has also been called Orb2 and Shk1 (Marcus et al., 1995; Verde et al., 1995). Pak1 function at the assembling CAR would likely require the presence of activated Cdc42, which binds to and activates Pak1. Cdc42 activity is downregulated during cytokinesis in budding yeast (Atkins et al., 2013), but may be active during fission yeast cytokinesis (Hercyk and Das, 2019). Using the well-established Cdc42 biosensor CRIB-3xGFP (Tatebe et al., 2008), we found that active GTP-Cdc42 localized to the cell middle and assembling CAR similar to Pak1 (Fig. 1C). Both active Cdc42 and Pak1 localized to the cell middle during the assembly and maturation phases of cytokinesis (Fig. 1C). We conclude that active Cdc42 and Pak1 enrich at the assembling CAR and might contribute to the early stages of cytokinesis, unlike other cell polarity kinases that reach the cell middle only at the later stages of cell division.

### Pak1 promotes CAR assembly and cytokinesis

Based on these localization results, we tested if Pak1 functions in CAR assembly. *pak1* is an essential gene, so we used two different mutant alleles: analog-sensitive *pak1-as* (M460A) and temperature-sensitive *orb2-34* (a mutant allele of *pak1*). Both mutants exhibit partial loss-of-function phenotypes including increased cell width under permissive conditions, consistent with severely reduced kinase activity *in vitro* (Fig. S1C-D; Loo and Balasubramanian, 2008; Das et al., 2012). We measured the time between SPB splitting and complete CAR assembly (Fig. S1E), and found that both *pak1* mutants took longer to assemble the CAR than wild type cells (Fig. 2A). CAR assembly was slowed even further by adding 3-Brb-PP1 inhibitor to *pak1-as* cells, which inhibits residual kinase activity in this mutant (Fig. S1F). In these mutants, longer CAR assembly was coupled with a shorter CAR maturation phase between assembly and constriction (Fig. S1G). Subsequent CAR constriction occurred with similar timing in the mutant and wild type cells (Fig. S1G). Thus, Pak1 localizes to the assembling CAR and is required for efficient CAR assembly.

**Figure 2:**
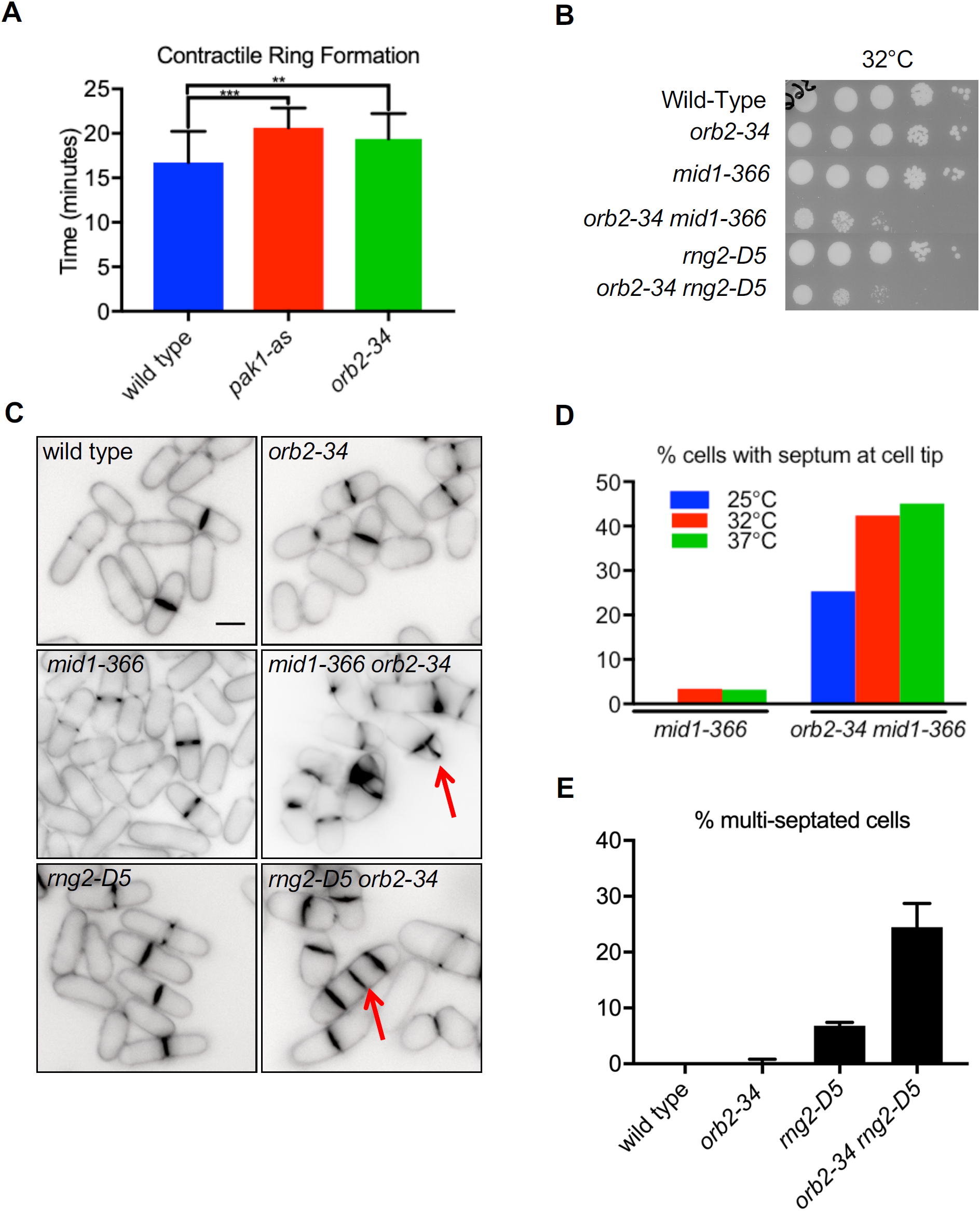
Cytokinesis defects in *pak1* mutants. **(A)** Quantification of CAR formation timing for the indicated strains (n ≥ 19 cells each). Values are mean ± SD. **p <0.01, ***p <0.001 **(B)** Serial-dilution growth assays for each indicated strains. **(C)** Representative images of strains grown at 25°C and stained with Blankophor. Red arrows indicate cytokinesis defects. Scale bar, 5 μm. **(D)** Percentage of cells with division septum at cell tip. n > 30 dividing cells for each strain and condition. **(E)** Percentage of cells with multiple septa (n > 60 cells each).

Genetic interactions provided additional support for the role of Pak1 in cytokinesis. We combined the *orb2-34* allele with *mid1-366* and *rng2-D5* temperature-sensitive mutants. Both Mid1/anillin and Rng2/IQGAP localize to cytokinesis nodes as well as the CAR and are critical for proper CAR formation (Eng et al., 1998; Wu et al., 2006, Laporte et al., 2011; Padmanabhan et al., 2011). We observed synthetic growth defects for both *orb2-34 mid1-366* and *orb2-34 rng2-D5* double mutants (Fig. 2B). These cells exhibited severe cytokinesis defects including misplaced, multiple, and/or disorganized division planes (Fig. 2C-E). We considered the possibility that these defects might be due to increased cell width in *orb2-34* mutants, as opposed to a more direct role in cytokinesis. To control for this possibility, we combined *mid1-366* and *rng2-D5* with *rga4Δ*, which exhibits increased cell width like *orb2-34* (Fig. S1C) (Das et al., 2007). No synthetic defects were observed with *rga4Δ* (Fig. S1H-I), indicating that *orb2-34* genetic interactions likely reflect a role for Pak1 in CAR assembly.

What proteins does Pak1 phosphorylate to promote cell division? One candidate substrate is the regulatory light chain of myosin II (Rlc1). Past work has shown that Pak1 phosphorylates Rlc1 at residues S35 and/or S36 (Loo and Balasubramanian, 2008). We performed *in vitro* kinase assays with purified proteins and found that Pak1 phosphorylates Rlc1-S36A, but not Rlc1-S35A or Rlc1-S35A,S36A (Fig. S2A-B). However, non-phosphorylatable *rlc1* mutant cells exhibited no defects in CAR assembly, cell shape, or genetic interactions with *mid1-366* (Fig. S2C-E). We conclude that Pak1 phosphorylates Rlc1, but this substrate does not explain the role of Pak1 in CAR assembly. Rather, these results suggest that additional cytokinesis proteins may be phosphorylated by Pak1 to promote cell division.

### Large-scale phosphoproteomic screen reveals novel Pak1 substrates

To identify Pak1 substrates in cell division, we quantitatively compared the phosphoproteome in wild type versus *pak1-as* cells using multiplexed tandem-mass-tag (TMT) labeling coupled with liquid chromatography and mass spectrometry. We tested 4 conditions: (1) *pak1-as* + methanol control, (2) *pak1-as* + 3-BrB-PP1, (3) wild type + methanol control, and (4) wild type + 3-BrB-PP1. By using 11-plex TMT labeling, we directly compared the abundance of tryptic phospho-peptides from 11 samples: the first 3 conditions in triplicate and the final condition in duplicate. By including replicates, we could calculate statistical significance of phosphorylation site changes between conditions. In total, we identified 11,292 phosphopeptides, 8,641 phosphosites and 1,848 phosphoproteins (Table S2). *pak1-as* cells exhibit cell polarity and cytokinesis defects in the absence of inhibitor due to dramatically reduced kinase activity, so we expected to find cell polarity and cytokinesis substrates when comparing *pak1-as* versus wild type, both lacking inhibitor. In this category, we identified 123 phosphopeptides reduced at least two-fold with a *p*-value of ≤ 0.05. This list included proteins that function in cell polarity (e.g. Scd1, Scd2, Tea3, and Rga4) and cytokinesis (e.g. Mid1, Cdc15, Cyk3, and Rng10) (Fig. 3A-B, Table S2).

**Figure 3:**
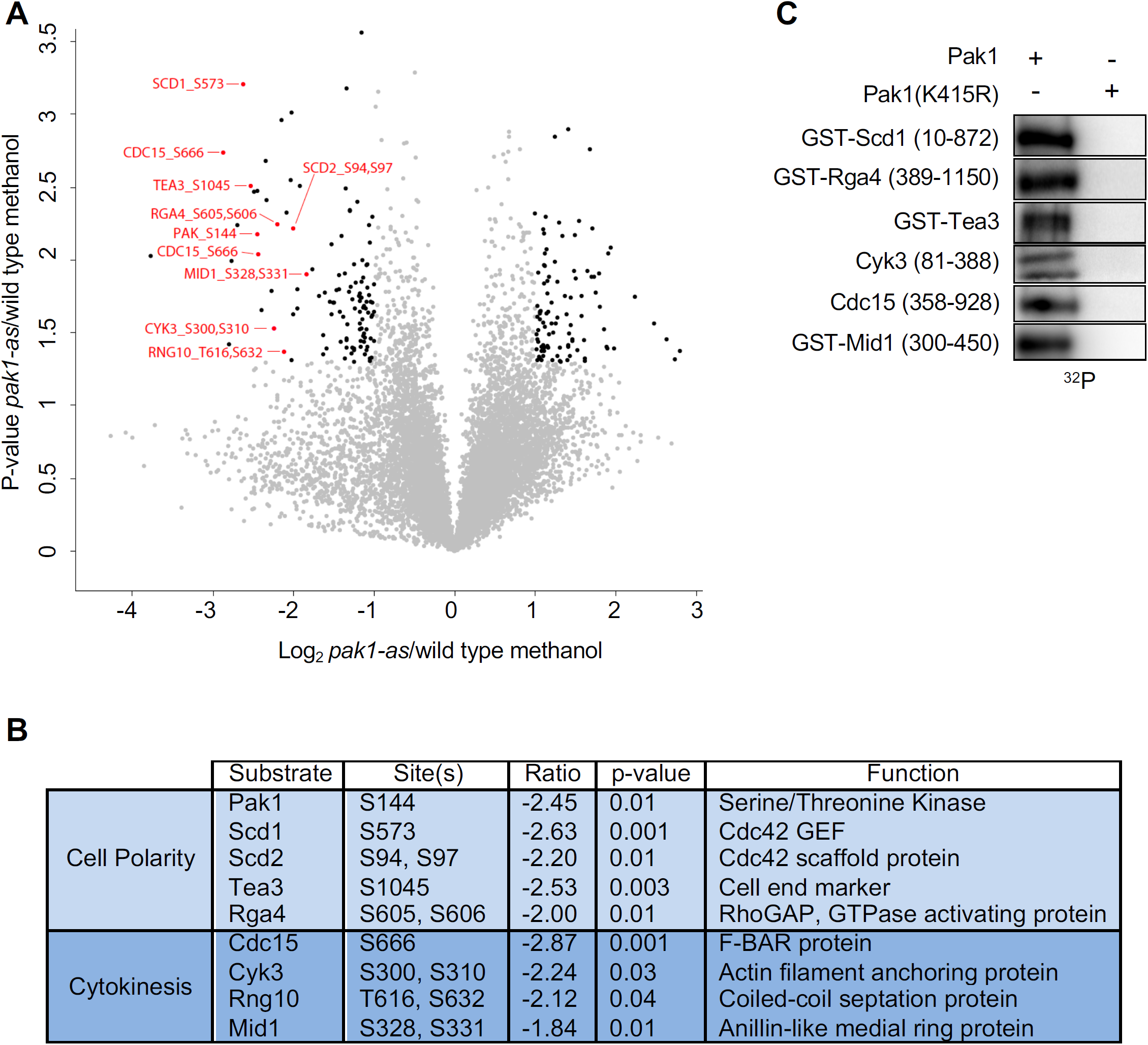
Large-scale phosphoproteomic screen reveals novel Pak1 substrates. **(A)** Volcano plot of phosphoproteomic data set comparing *pak1-as* versus wild type cells. Each point represents an identified phosphopeptide. Protein name and modified residues are shown. **(B)**List of putative Pak1 substrates involved in cell polarity and cytokinesis. **(C)** *In vitro* kinase assays using purified proteins and γ-^32^P-ATP.

We verified several targets as direct substrates using *in vitro* kinase assays with purified proteins and γ-^32^P-ATP. We purified full-length Tea3 and fragments of Scd1, Rga4, Cyk3, Cdc15 and Mid1 that contained residues phosphorylated in a Pak1-dependent manner in cells (Fig. S2F). These proteins were phosphorylated by Pak1 but not by kinase-dead Pak1(K415R) *in vitro*, showing that they are direct Pak1 substrates (Fig. 3C).

### Pak1 kinase activity regulates cellular localization of Cdc15 and Mid1

We focused on the new Pak1 substrates Cdc15 and Mid1 due to their roles in cytokinesis. Cdc15 is a membrane-binding phosphoprotein that connects multiple cytokinesis factors to the CAR (Roberts-Galbraith et al., 2010; Martín-García et al., 2014; Kettenbach et al., 2015; Ren et al., 2015; Willet et al., 2015; Lee et al., 2018, Bhattacharjee et al., 2020) and exhibits synthetic genetic defects with *pak1* mutations (Fig. 4A-B). During cytokinesis, Cdc15 completely shifts localization from cell tips to the CAR. This shift was defective in *pak1-as*, leading cells to undergo cytokinesis with residual Cdc15 at cell tips (Fig. 4C-E). This result suggests that Pak1 alters Cdc15 dynamics to promote its redistribution to the cell middle. Consistent with this notion, we observed abnormally stable Cdc15 puncta at the tips of *pak1-as* but not wild type cells (Fig. S3A-B). Taken together, these results show that Pak1 directly phosphorylates Cdc15 to promote its concentration in the CAR for cytokinesis.

**Figure 4:**
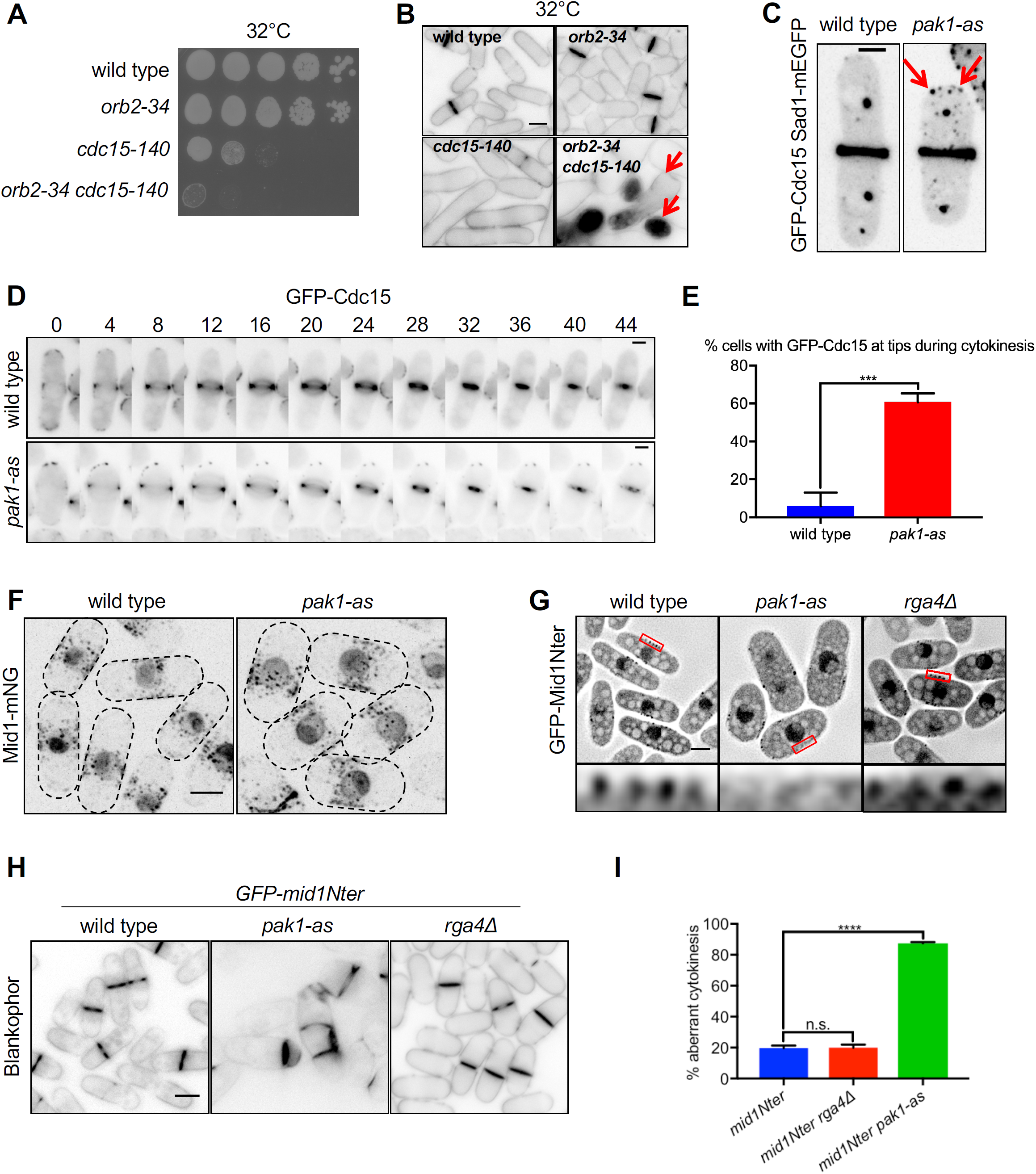
Pak1 promotes recruitment of its substrates Mid1 and Cdc15 to CAR. **(A)** Serial-dilution growth assays for each indicated strain. **(B)** Representative images of cells stained with Blankophor. **(C)** Images of GFP-Cdc15 Sad1-mEGFP during cytokinesis. Arrows indicate Cdc15 localization at cell tips during cytokinesis in *pak1-as* cells. Images are maximum intensity projections. Scale bar, 2 μm. **(D)** Representative time-lapse montage of GFP-Cdc15 during cytokinesis. Note Cdc15 localization at cell tips in *pak1-as*. Images are single focal planes. Scale bar, 2 μm. **(E)** Quantification of cells with GFP-Cdc15 at cell tips during cytokinesis. Values are the mean ± SD from three biological replicates (n > 50 cells each), ***p <0.001. **(F)** Localization of Mid1-mNG in the indicated strains. Images are maximum intensity projections. Dotted lines outline cell boundaries. Scale bar, 5 μm. **(G)** Localization of GFP-Mid1Nter. Insets are enlarged views of boxed region. Scale bar, 3 μm. **(H)** Images of GFP-Mid1Nter cells stained with Blankophor. Scale bar, 5 μm. **(I)** Quantification of aberrant cytokinesis in Blankophor-stained cells. Values are mean ± SD from three biological replicates (n ≥ 75 cells each), ****p <0.0001.

Next, we investigated the anillin-related protein Mid1, which localizes to CAR node precursors and the mature CAR, and is required to position the CAR in the cell middle (Celton-Morizur et al., 2006; Padte et al., 2006; Almonacid et al., 2009; Almonacid et al., 2011; Chang et al., 1996; Paoletti and Chang, 2000). The Mid1 N-terminus (Mid1Nter, residues 1-506) localizes to nodes and the CAR, and is both necessary and sufficient for Mid1 function in cytokinesis (Guzman-Vendrell et al., 2013). The Mid1 C-terminus contains a membrane-binding amphipathic helix that reinforces Mid1 localization (Celton-Morizur et al., 2004). We did not observe obvious defects in the localization of Mid1-mNG in *pak1-as* cells (Fig. 4F), so we focused on the Mid1Nter construct that contains Pak1 sites identified both *in vivo* and *in vitro*. Interestingly, these phosphorylation sites cluster in binding domains for Cdr2 and Gef2, two cortical node proteins that are essential for Mid1Nter localization and function (Almonacid et al., 2009, Ye et al., 2012). Consistent with past work, GFP-Mid1Nter localized to cortical nodes in the cell middle and supported proper positioning of the cell division plane (Guzman-Vendrell et al., 2013). However, both the localization and function of Mid1Nter were disrupted in *pak1-as* cells. GFP-Mid1Nter still localized to the cell cortex in *pak1-as* cells, but not in nodes that concentrated in the cell middle (Fig. 4G). This mislocalization correlated with dramatic cytokinesis defects, as *GFP-mid1Nter pak1-as* cells exhibited a range of division plane defects including misplaced, split, and tilted septa (Fig. 4H-I). These defects were not simply due to the increased width of *pak1-as* cells, as we did not observe similar phenotypes or localization defects for *GFP-mid1Nter rga4Δ* cells (Fig. 4G-I). These results suggest that Pak1 phosphorylates the N-terminus of Mid1 to promote proper cytokinesis.

### Pak1 promotes Mid1 localization to cortical nodes

To gain more insight into Pak1 regulation of Mid1, we mutated 9 serines phosphorylated by Pak1 *in vitro* and/or in cells to alanine (Fig. 5A and Table S2). The resulting Mid1(9A) mutant almost completely abolished phosphorylation by Pak1 *in vitro* (Fig. 5B). In cells, the GFP-Mid1Nter(9A) mutant failed to localize at nodes and exhibited severe cytokinesis defects (Fig. 5C-D). These defects phenocopy combining *pak1-as* with GFP-Mid1Nter. Interestingly, these defects were not due to loss of nodes themselves, because the core node protein Cdr2 still localized properly in the GFP-Mid1Nter(9A) mutant (Fig. S3C). These results suggest that Pak1 phosphorylates Mid1 to promote its recruitment to Cdr2 nodes, consistent with these phosphorylation sites being found within binding domains for Cdr2 and Gef2.

**Figure 5:**
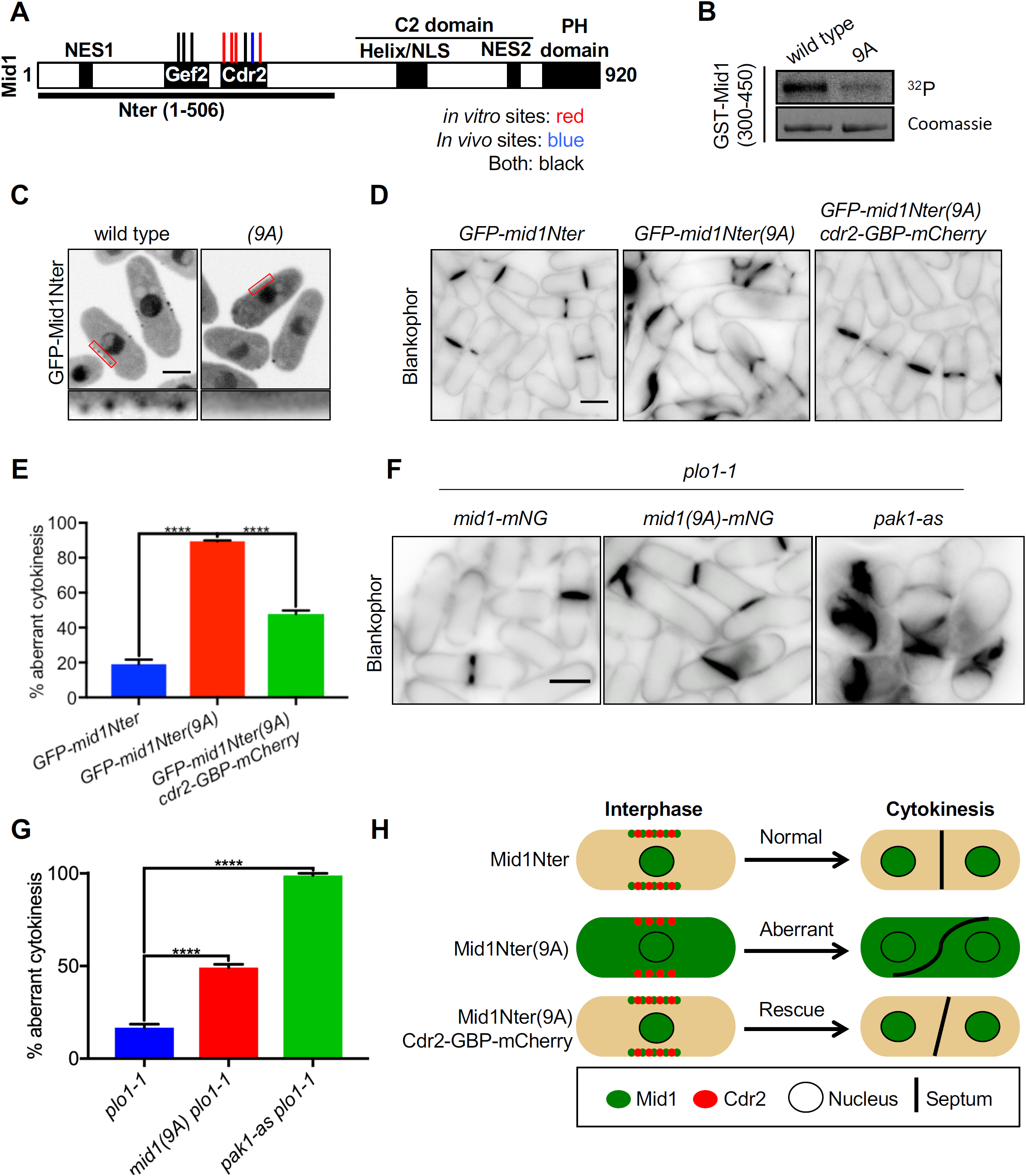
Pak1 phosphorylates the N-terminus of Mid1 to ensure proper cytokinesis. **(A)** Schematic of Mid1 domain layout. Pak1 phosphorylation sites are indicated by red, blue or black lines. **(B)** *In vitro* kinase assay with purified proteins and γ-^32^P-ATP. **(C)** Localization of GFP-Mid1Nter. Single focal plane images are shown. Insets are enlarged views of boxed region. Scale bar, 3 μm. **(D)** Images of GFP-Mid1Nter cells stained with Blankophor. Scale bar, 3 μm. **(E)** Quantification of aberrant cytokinesis measured from Blankophor stained cells. Values are the mean ± SD from three biological replicates (n ≥ 75 cells each), ****p <0.0001. **(F)** Images of indicated strains stained with Blankophor. Scale bar, 5 µm. **(G)** Quantification of aberrant cytokinesis measured from Blankophor stained cells. Values are the mean ± SD from three biological replicates (n ≥ 75 cells each), ****p <0.0001. **(H)** Model for Pak1 regulation of Mid1.

If Mid1Nter(9A) cytokinesis defects are due simply to mislocalization as opposed to other aspects of function, then synthetic recruitment of Mid1Nter(9A) back to nodes should suppress the cytokinesis defects. We tested this prediction using the GFP binding peptide (GBP) system. In *GFP-mid1Nter(9A) cdr2-GBP-mCherry* cells, Mid1Nter(9A) localization was restored to cortical nodes (Fig. S3D). More importantly, this synthetic retargeting largely suppressed the cytokinesis defects and restored proper cell division to this mutant (Fig. 5D-E, Fig. S3E-F). These results show that Pak1 phosphorylates the N-terminus of Mid1 to promote node localization for proper cytokinesis.

Finally, we tested the effects of Pak1 regulation by generating full-length Mid1(9A). We did not observe cytokinesis defects in *mid1(9A)* mutants (Fig. S3G-H) but tested genetic interactions with mutants in Polo kinase Plo1. Plo1 phosphorylates the Mid1 N-terminus at distinct sites from Pak1 to promote Mid1 nuclear export and downstream function in recruiting cytokinesis proteins such as Rng2 to nodes (Bähler et al., 1998, Alamonacid et al., 2011). We discovered synthetic defects in cytokinesis when combining *mid1(9A)* with *plo1-1* (Fig. 5F-G). Interestingly, *plo1-1* exhibits similar synthetic defects with other mutants that impair Mid1 anchoring to nodes including *cdr2Δ* and *gef2Δ* (Almonacid et al., 2009, Ye et al., 2012). Similar to *mid1(9A)*, combining *pak1-as* with *plo1-1* also led to severe cytokinesis defects (Fig. 5F-G). Combined with previous work, our results suggest that both Pak1 and Plo1 kinases phosphorylate Mid1 to promote distinct steps in its regulatory mechanism. Both kinases phosphorylate the Mid1 N-terminus but at distinct residues and domains. Plo1 phosphorylates Mid1 to promote its export from the nucleus and its recruitment of cytokinesis factors IQGAP and myosin II (Bähler et al., 1998, Almonacid et al., 2011). In contrast, Pak1 phosphorylates Mid1 to promote its association with Cdr2 nodes (Fig. 5H). Mutations targeting either step alone do not generate strong defects, but impairing both mechanisms leads to strong cytokinesis defects. This two-pronged regulatory network ensures that Mid1 recruits cytokinesis proteins in the correct time (Plo1) and place (Pak1).

### Conclusions

Our search for cell polarity kinases that function in cytokinesis revealed recruitment of Pak1 and its upstream activator Cdc42 to the assembling CAR for cytokinesis. We found new functions for Pak1 in cytokinesis and used large-scale phosphoproteomics to identify new substrates that are directly phosphorylated by Pak1 *in vitro*. We showed that Mid1 is a critical substrate for Pak1 function in cytokinesis. Pak1 phosphorylates the N-terminus of Mid1 to promote its association with cortical nodes. We note that this regulatory mechanism could involve the recently described phase separation of this domain (Chatterjee and Pollard, 2019), a possibility that could be examined in future work. Pak1 regulation of Mid1 acts in conjunction with other kinases including Plo1 and Sid2 (Almonacid et al., 2011, Willet et al., 2019). Each kinase regulates distinct aspects of Mid1 function, suggesting a coordinated regulatory network with implications for cytokinetic scaffolds such as anillin in other cell types. The full role of Pak1 in cytokinesis likely involves proteins beyond Mid1, consistent with our identification of Cdc15, Cyk3, and Rng10 as new substrates for Pak1. We note that Cdc15 and Cyk3 are also phosphorylated by the cell polarity kinases Pom1 and Kin1 (Kettenbach et al., 2015; Lee et al., 2018; Bhattacharjee et al., 2020), and Pom1 further phosphorylates the Rng10 ligand Rga7 (Kettenbach et al., 2015). Put together, these connections reveal a strong and growing role for cell polarity signaling in the regulation of CAR proteins for cytokinesis.

## Figure legends

**Supplemental Figure S1:**
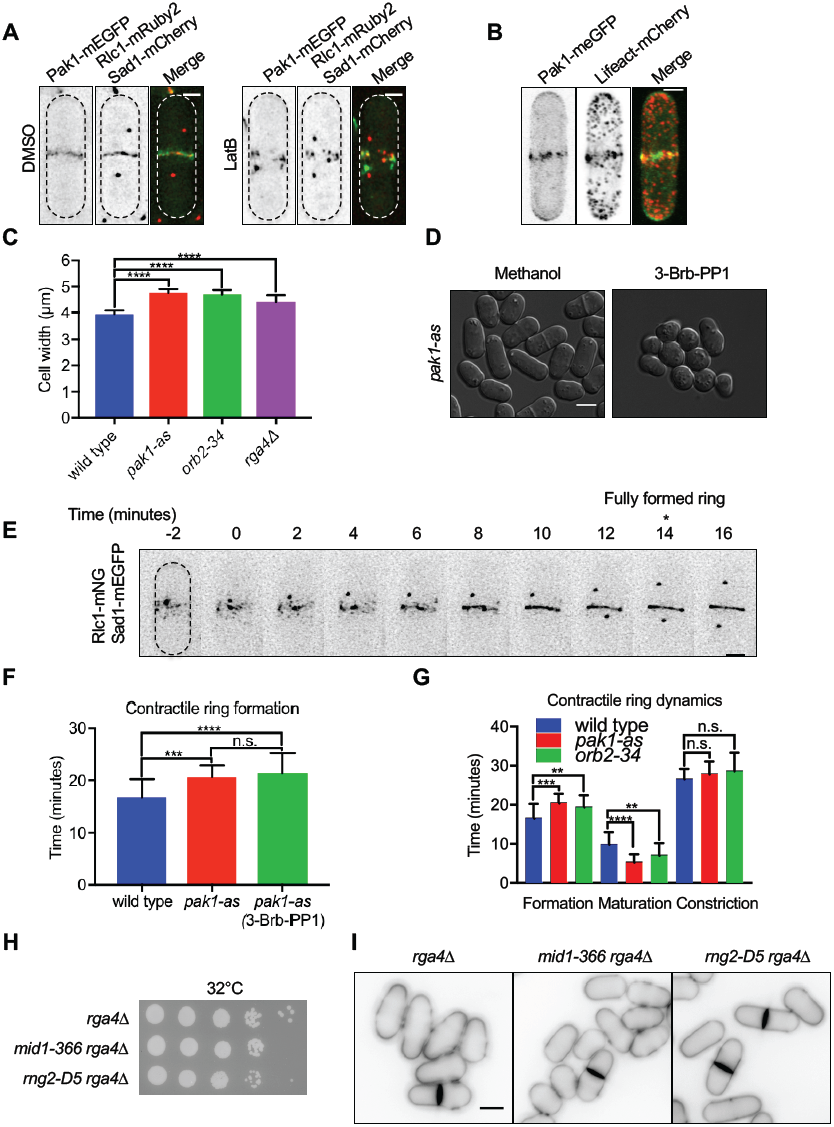
Additional characterization of *pak1* mutant phenotypes. **(A)** Localization of indicated proteins in cells treated with DMSO or 100μM Latrunculin B. Images are maximum intensity projections. Dotted lines outline cell boundaries. **(B)** Colocalization of Pak1-mEGFP and LifeAct-mCherry by Airyscan confocal microscopy. Images are maximum intensity projections. **(C)** Quantification of cell width for indicated strains (n ≥ 50 cells each). Values are the mean ± SD, **** p <0.0001. **(D)** DIC images of *pak1-as* cells treated overnight at 25°C with either 30μM 3-Brb-PP1 or methanol. **(E)** Time lapse montage of Rlc1-mNG Sad1-mEGFP localization during cytokinesis. SPB splitting is time zero for monitoring CAR formation. **(F)** Quantification of CAR formation for each indicated strain (n ≥ 18 cells each). For *pak1-as* 3-Brb-PP1, cells were briefly incubated with 30μM 3-Brb-PP1 and then immediately imaged on YE4S agarose pads containing 30μM 3-Brb-PP1. Values are mean ± SD. ***p <0.001, ****p <0.0001. **(G)** Quantification of CAR dynamics in the indicated strains (n ≥ 18 cells each). Values are mean ± SD. **p <0.01, ***p <0.001, ****p <0.0001. **(H)** Serial-dilution growth assays for each indicated strain. Strains were grown at 32°C for 3-4 days on YE4S media. **(I)** Images of the indicated *rga4Δ* strains stained with Blankophor. Scale bars (A-B, E) 2 μm; (D, I) 5 μm.

**Supplemental Figure S2:**
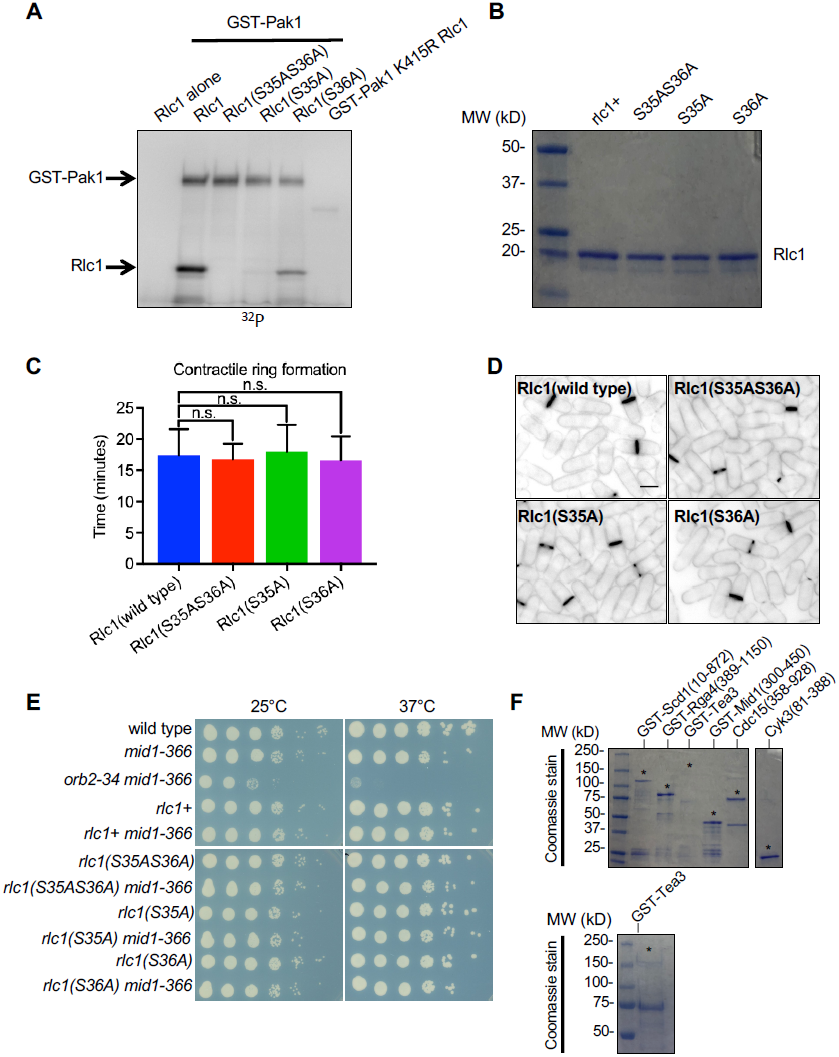
*pak1* cytokinesis defects are independent of Rlc1 phosphorylation. **(A)** *In vitro* kinase assay using full-length Pak1 and Rlc1 purified from bacteria. The indicated proteins were mixed with γ-^32^P-ATP, separated by SDS-PAGE and visualized by autoradiography. **(B)** Purified proteins used for *in vitro* kinase assays in Fig. S2A. Proteins were separated by SDS-PAGE followed by Coomassie blue staining. **(C)** Quantification of CAR formation timing for each indicated strains (n = 20 cells each). Values are mean ± SD. **(D)** Images of indicated strains stained with Blankophor. Scale bar, 5 μm. **(E)** Serial-dilution growth assays. The indicated strains were grown at either 25°C or 37°C for 3-4 days on YE4S plates. **(F)** Purified proteins used for *in vitro* kinase assays in Fig. 3C. Proteins were separated by SDS-PAGE followed by Coomassie blue staining. Proteins were loaded at concentrations used for *in vitro* kinase assays, and asterisks denote relevant protein band. Lower panel shows separate gel with higher loading volume for GST-Tea3 to confirm band size.

**Supplemental Figure S3:**
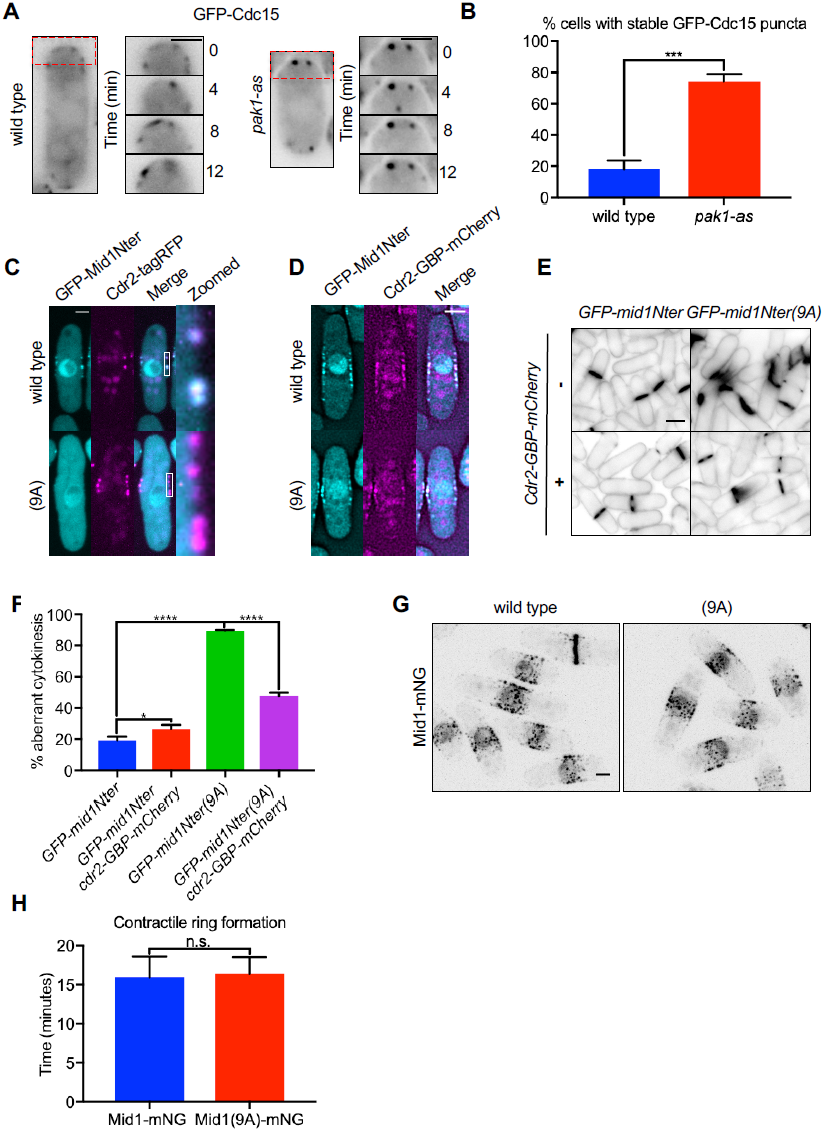
Characterization of Mid1 and Cdc15 in *pak1* mutants. **(A)** Single focal plane images of GFP-Cdc15 from time-lapse microscopy. Whole cell image represents time point zero. Insets are cell tip images over time from the boxed region. Scale bars, 2 μm. **(B)** Quantification of cells with stable GFP-Cdc15 puncta at cell tips. Values are the mean ± SD from three biological replicates (n > 50 cells each), ***p <0.001. **(C)** Colocalization of GFP-Mid1Nter and Cdr2-tagRFP. Single focal plane spinning disc confocal images are shown. Insets are enlarged views of boxed region. Scale bar, 2 μm. **(D)** Colocalization of GFP-Mid1Nter and Cdr2-GBP-mCherry. Single focal plane deconvolved widefield images are shown. **(E)** Blankophor images of indicated strains. Scale bar, 5 μm. **(F)** Quantification of aberrant cytokinesis judged from Blankophor stained cells. Values are the mean ± SD from three biological replicates (n ≥ 75 cells each), *p <0.05, ****p <0.0001. **(G)** Localization of Mid1-mNG in the indicated strains. Images are maximum intensity projections acquired by spinning disc confocal microscopy. Scale bar, 3 μm. **(H)** Quantification of CAR dynamics in the indicated strains (n = 20 cells each). Values are mean ± SD.

**Table S1: List of yeast strains and plasmids used in this study**

**Table S2: Phosphoproteomic data set**

## Materials and methods

### Strain construction and media

Standard *S. pombe* media and methods were used (Moreno et al., 1991). Strains used in this study are listed in Table S1. PCR-based homologous recombination was performed for chromosomal tagging and deletion (Bähler et al., 1998). The *pak1-as* (M460A) strain was generated and shared by Juraj Gregan (Cipak et al., 2011). The non-phosphorylatable *rlc1* sequences were generated by site-directed mutagenesis using QuikChange II mutagenesis (Stratagene) according to the manufacturers protocol, and integrated into the *leu1* locus of JM837 (*leu1-32*) using pJK148. Strains with *rlc1* sequences integrated at the *leu1* locus were crossed to a *rlc1Δ* strain to eliminate the copy at the endogenous locus. All plasmids were sequenced by Sanger Sequencing for verification. For growth assays, cells were spotted by 10-fold serial dilutions on YE4S plates and were incubated at 25°C, 32°C or 37°C for 3-4 days before scanning. *Pmid1-mid1-mNeonGreen-Tadh1* was generated by PCR from genomic DNA of strain JM5288 and inserted into pJK210 by restriction digest and ligation. *Pmid1-GFP-mid1Nter-Tnmt1* was generated by PCR from genomic DNA of strain JM1493 and inserted into pJK210 by restriction digest and ligation. The *mid1(9A)* mutant allele was synthesized as a gBlocks Gene Fragment (Integrated DNA Technologies), and then inserted into the PCR-linearized fragment from pJK210*-Pmid1-mid1-mNeonGreen-Tadh1* or pJK210*-Pmid1-GFP-mid1Nter-Tnmt1* plasmid by repliQa HiFi Assembly Mix (Quantabio) according to the manufacturer’s protocol. For the 9A mutant, the following residues were mutated to alanine: S328, S329, S331, S403, S416, S418, S422, S425, and S432. The wild type and non-phosphorylatable *mid1* alleles were integrated into the *ura4* locus of a *mid1Δ* strain.

### Large-Scale Phosphoproteomic Screen

The strains used for phosphoproteomic screen were JM366 (972) and JM4787 (*pak1::ClonNatR pak1-as(M460A)-HphR*). Cells were grown in YE4S at 32°C for at least 8 generations and treated for 15 minutes with 30μM 3-Brb-PP1 (3-[(3-Bromophenyl)methyl]-1- (1,1-dimethylethyl)-1H-pyrazolo[3,4-d]pyrimidin-4-amine (Abcam), while control culture was treated with equal volume of methanol. Each culture volume was 1 liter YE4S. The cultures were then harvested by 10-minute centrifugation at 8,000 g at 4°C, washed once with 200 mL ice-cold 1x PBS, and then centrifuged again. The remaining cell pellet was weighed and resuspended in a 1:1 ratio with ice-cold 1x PBS with a protease inhibitor tablet (Roche Life Sciences) and 1mM PMSF. The cells were then lysed by two minutes of grinding in a prechilled coffee bean grinder; lysis efficiency was ∼80% as judged by microscopy.

Yeast powder was lysed in ice-cold lysis buffer ((8 M urea, 25 mM Tris-HCl pH 8.6, 150 mM NaCl, phosphatase inhibitors (2.5 mM beta-glycerophosphate, 1 mM sodium fluoride, 1 mM sodium orthovanadate, 1 mM sodium molybdate) and protease inhibitors (1 mini-Complete EDTA-free tablet per 10 ml lysis buffer; Roche Life Sciences)) and sonicated three times for 15 seconds each with intermittent cooling on ice. Lysates were centrifuged at 15,000 x g for 30 minutes at 4°C. Supernatants were transferred to a new tube and the protein concentration was determined using a BCA assay (Pierce/ThermoFisher Scientific). For reduction, DTT was added to the lysates to a final concentration of 5 mM and incubated for 30 min at 55°C. Afterwards, lysates were cooled to room temperate and alkylated with 15 mM iodoacetamide at room temperature for 45 min. The alkylation was then quenched by the addition of an additional 5 mM DTT. After 6-fold dilution with 25 mM Tris-HCl pH 8, the samples were digested overnight at 37°C with 1:100 (w/w) trypsin. The next day, the digest was stopped by the addition of 0.25% TFA (final v/v), centrifuged at 3500 x g for 30 minutes at room temperature to pellet precipitated lipids, and peptides were desalted on a 500 mg (sorbent weight) SPE C18 cartridge (Grace-Davidson). Peptides were lyophilized and stored at −80°C until further use.

Phosphopeptide purification was performed as previously described (Kettenbach and Gerber, 2011). Briefly, peptides were resuspended in 1.5 M lactic acid in 50% ACN (“binding solution”). Titanium dioxide microspheres were added and vortexed by affixing to the top of a vortex mixer on the highest speed setting at room temperature for 1 hour. Afterwards, microspheres were washed twice with binding solution and three times with 50% ACN / 0.1% TFA. Peptides were eluted twice with 50 mM KH_2_PO_4_ (adjusted to pH 10 with ammonium hydroxide). Peptide elutions were combined, quenched with 50% ACN / 5% formic acid, dried and desalted on a µHLB OASIS C_18_ desalting plate (Waters). Phosphopeptide enrichment was repeated once.

Phosphopeptides were resuspended in 133 mM HEPES (SIGMA) pH 8.5 and 20% acetonitrile (ACN) (Burdick & Jackson). Peptides were transferred to dried, individual TMT reagent (ThermoFisher Scientific), and vortex and mix reagent and peptides. After 1 hr at room temperature, each reaction was quenched with 3 µl of 500 mM ammonium bicarbonate solution for 10 minutes, mixed, diluted 3-fold with 0.1% TFA in water, and desalted using C_18_ solid phase extraction cartridges (ThermoFisher Scientific). The desalted multiplex was dried by vacuum centrifugation and separated by offline Pentafluorophenyl (PFP)-based reversed phase HPLC fractionation was performed as previously described (Grassetti, Hards et al., 2017).

TMT-labeled samples were analyzed on a Orbitrap Fusion (Senko, Remes et al., 2013) mass spectrometer (ThermoScientific) equipped with an Easy-nLC 1000 (ThermoScientific). Peptides were resuspended in 8% methanol / 1% formic acid across a column (45 cm length, 100 μm inner diameter, ReproSil, C_18_ AQ 1.8 μm 120 Å pore) pulled in-house across a 2 hrs gradient from 8% acetonitrile/0.0625% formic acid to 37% acetonitrile/0.0625% formic acid. The Orbitrap Fusion was operated in data-dependent, SPS-MS3 quantification mode (Ting, Rad et al., 2011, McAlister, Nusinow et al., 2014) wherein an Orbitrap MS1 scan was taken (scan range = 350 – 1500 m/z, R = 120K, AGC target = 2.5e5, max ion injection time = 100ms), followed by ion trap MS2 scans on the most abundant precursors for 4 seconds (max speed mode, quadrupole isolation = 0.6 m/z, AGC target = 4e3, scan rate = rapid, max ion injection time = 60ms, minimum MS1 scan signal = 5e5 normalized units, charge states = 2, 3 and 4 included, CID collision energy = 33%) and Orbitrap MS3 scans for quantification (R = 15K, AGC target = 2e4, max ion injection time = 125ms, HCD collision energy = 48%, scan range = 120 – 140 m/z, synchronous precursors selected = 10). The raw data files were searched using COMET with a static mass of 229.162932 on peptide N-termini and lysines and 57.02146 Da on cysteines, and a variable mass of 15.99491 Da on methionines and 79.96633 Da on serines, threonines and tyrosine against the target-decoy version of the *S. pombe* FASTA database (UniProt; www.uniprot.org) and filtered to a <1% FDR at the peptide level. Quantification of LC-MS/MS spectra was performed using in house developed software. Phosphopeptide intensities were adjusted based on total TMT reporter ion intensity in each channel and log_2_ transformed. P-values were calculated using a two tailed Student’s t-test assuming unequal variance.

For volcano plot in Figure 3A, gray points are not significantly different between *pak1-as* and wild type cells treated with methanol. Black points have ≥ 2-fold decrease in abundance in *pak1-as* vs. wild type cells with a p-value of ≤ 0.05. Red points are significant phosphopeptides identified that have roles in polarized growth and cytokinesis. Protein name and modified residues are shown. Note that 5 outliers were removed from the graph for clarity of presentation. They are Pma1 S911:T912, Plc1 Y296:S297, Leu3 S355, Igo1 S128 and Rct1 S206. These phosphopeptides are included in the full data set found in Table S2. For Fig. 3B, each phosphopeptide’s abundance ratio between *pak1-as* and wild type (both in methanol) was log_2_-transformed.

### Protein purification *and in vitro* kinase assays

Full length wild type and catalytically inactive Pak1, wild type and mutant Rlc1, Tea3, and fragments of Mid1 (aa 300-450), Mid1(9A) (aa 300-450), Scd1 (aa 10-872) and Rga4 (aa 389-1150) were cloned into pGEX6P1 (GST tag) vector (GE Healthcare), expressed in BL21 (DE3 or Rosetta) E. coli strains, and purified with glutathione agarose resin (Sigma-Aldrich) as previously described (Kettenbach et al., 2015). Purified proteins were released from resin by overnight incubation with 3C protease at 4°C or by elution with glutathione. Fragments of Cyk3 (aa 81-388) and Cdc15 (aa 358-928) were purified as previously described (Lee et al., 2018).

*In vitro* kinase assays were performed by incubating purified proteins in kinase assay buffer (50 mM Tris-HCL pH 7.5, 150 mM NaCl, 10 mM MgCl_2_, 1 mM MnCl_2_) supplemented with 10 μM ATP and 2 μCi γ-^32^P-ATP (blu002z250uc, Perkin Elmer) in 15μL reactions. Reactions were incubated at 30°C and stopped after 30 minutes by adding SDS-PAGE sample buffer (65 mM Tris, pH 6.8, 3% SDS, 10% glycerol, 10% 2-mercaptoethanol) and boiling. Gels were dried for one hour and the signal of γ-^32^P-ATP was detected via a PhosphorImager scanned by Typhoon 8600 Variable Mode Imager (GE Healthcare). For Fig. 5B, reactions were incubated at 30°C and stopped after 5 minutes by adding SDS-PAGE sample buffer (65 mM Tris, pH 6.8, 3% SDS, 10% glycerol, 10% 2-mercaptoethanol) and boiling. Gels were dried for one hour and the signal of γ-^32^P-ATP was detected via film exposure for 1 hour in a dark room. In all cases, bands in the figures correspond to the molecular weight where coomassie-stained protein migrates.

For identification of Pak1 phosphorylation sites on Mid1 *in vitro*, purified full length wild type and catalytically inactive Pak1 was incubated with GST-Mid1 (300-450) for 1 hour at 30°C in kinase assay buffer supplemented with 10 μM ATP. Reactions were stopped by addition of buffer (1% SDS, 15% glycerol, 50mM Tris-HCL, pH 8.7, 150 mM NaCl) and boiled for 5 min at 80°C, reduced with 5mM DTT, and alkylated with 15mM iodoacetamide (Sigma-Aldrich) for 1 hour in dark prior to boiling in SDS-PAGE sample buffer for 5 min at 99°C. Reactions were then separated by SDS-PAGE. Gel bands were excised, destained, trypsin-digested and peptides were extracted and analyzed by LC-MS/MS (Table S2) (Opalko et al., 2020).

### Widefield microscopy and analysis

Cells were imaged in either EMM4S or YE4S at either 25°C, 32°C or 37°C using a DeltaVision imaging system (Applied Precision Ltd.) composed of an IX-inverted widefield microscope (Olympus) with a 100x or 60x oil objective, a CoolSnap HQ2 camera (Photometrics), and an Insight solid-state illumination unit (Applied Precision Ltd.). Images shown as Z stacks were acquired and processed by iterative deconvolution using SoftWoRx software (Applied Precision Ltd.). Single channel fluorescence images are shown in inverted look-up table (LUT) except for Fig. S3D. For Fig. 1A, images are shown as maximum intensity projections. For Figs. 2A and S1E-G, cells were placed onto agarose pads containing the same growth medium and 2% agarose. Z-stacks of cells were imaged every two minutes and analyzed as maximum intensity projections. For Fig. 4D, the medial focal plane of cells were imaged on YE4S agarose pads every two minutes. We counted cells with detectable GFP-Cdc15 signal at cell tips upon CAR constriction. Figs. 4G and S3D are displayed as a deconvolved single focal plane to resolve cortical nodes. All other single focal plane images are non-deconvolved. The criteria for aberrant cytokinesis analysis in Figs. 4I, 5D, and S3F were tilted septa, multiple septa, tip septa and split septa. For Fig. 5G, cells were grown overnight at 32°C in EMM4S in a shaking waterbath, stained with blankophor before imaging. The criteria for aberrant cytokinesis analysis in Fig. 5G were oblique septa, multiple septa, tip septa, split septa and aberrant cell wall deposition during division. For Fig. S1C, cell width was measured by imaging cells with stained Blankophor and drawing a line across the division plane of septating cells. For Fig. S3A, cells with GFP-Cdc15 puncta that persisted for at least twelve minutes were marked as cells with stable GFP-Cdc15 puncta. All image analysis was performed on ImageJ (National Institutes of Health). Statistical differences were assessed by either One-Way ANOVA or Welch’s t test using GraphPad.

### Spinning disc microscopy and analysis

Images for Figs. 1C, 4C, 4F, 5C, S2C and S3C, G, and H were taken with a spinning-disc confocal system (Micro Video Instruments, Avon, MA) featuring a Nikon Eclipse Ti base equipped with an Andor CSU-W1 two-camera spinning disc module, dual Zyla sCMOS cameras (Andor, South Windsor, CT) an Andor ILE laser module, and a Nikon 100X Plan Apo λ 1.45 oil immerision objective. Figs. 1C, 4C, 4F, and S3G are displayed as maximum intensity projections. Cells for timelapse imaging in for Figs. S2C and S3H were imaged on YE4S agarose pads. Z-stacks were acquired every two minutes and analyzed as maximum intensity projections. For Figs. 5C and S3C, medial focal plane images were acquired. Statistical differences were assessed by One-Way ANOVA using GraphPad.

### Airyscan super resolution microscopy and analysis

Cells were imaged using a Zeiss Airyscan microscope (Figs. 1B and S1B), composed of a Zeiss LSM-880 laser scanning confocal microscope (Zeiss, Oberkochen, Germany) equipped with 100X alpha Plan-Apochromat/NA 1.46 Oil DIC M27 Elyra objective, Airyscan super-resolution module and GaAsp Detectors, and Zen Blue acquisition software using the Super-resolution mode. Z-volumes of 32 slices with 0.17μm spacing through the cell. Airyscan images were processed in Zeiss Zen Blue software and presented as maximum intensity Airyscan reconstructed stacks. Images in Fig. 1B are Airyscan reconstructed maximum intensity projection Z-stacks of 34 sections with 0.17 μm spacing. Images in Fig. S1A are Airyscan reconstructed maximum intensity projection Z-stacks of 28 sections with 0.17 μm spacing. The delay in acquisition between the two channels in these experiments was 4 seconds.

### Statistical analyses

Welch’s T-test was used to assess differences for Figs. 4E, S3B and S3H. One-Way ANOVA followed by Dunnett’s multiple comparison test was used to assess differences for Figs. 2A, 4I, 5G, S1C, S1G and S2C. One-Way ANOVA followed by Tukey’s multiple comparison test was used to assess differences for Figs. 5E, S1F and S3F. All statistical analysis was done using GraphPad Prism 7.

## Acknowledgements

We thank members of the Moseley laboratory for comments on the manuscript, as well as L. Myers at Dartmouth for help with *in vitro* kinase assays, A. Lavanway for assistance with microscopy, and M. Adamo for help with mass spectrometry data. We thank the Biomolecular Targeting Core (BioMT) (P20-GM113132) and the Imaging Facility at Dartmouth for use of equipment; and J.-Q. Wu, A. Paoletti, J. Gregan for sharing strains.

This work was supported by grants from the American Cancer Society (RSG-15-140-01) and the National Institute of General Medical Sciences (R01GM099774 and R01GM133856) to J.B.M, and by grants from NIH/NIGMS (R35GM119455, P20GM113132) to A.N.K., and by the NCCC support grant from NIH/NCI (P30CA023108). The Orbitrap Fusion Tribrid mass spectrometer was acquired with support from NIH (S10-OD016212).

The authors declare no competing financial interests.

## Author contributions

J.B. Moseley and J.O. Magliozzi conceived of the study; J. Sears L. Cressey, and A.N. Kettenbach performed and analyzed mass spectrometry experiments; J.O. Magliozzi, M. Brady, and H.E. Opalko performed all other experiments; all authors analyzed the data; J.B. Moseley and J.O. Magliozzi wrote the manuscript; all authors edited the manuscript.

## References

Almonacid, M., Celton-Morizur, S., Jakubowski, J.L., Dingli, F., Loew, D., Mayeux, A., Chen, J.-S., Gould, K.L., Clifford, D.M., and Paoletti, A. (2011). Temporal control of contractile ring assembly by Plo1 regulation of myosin II recruitment by Mid1/anillin. Curr. Biol. 21, 473–479.

Almonacid, M., Moseley, J.B., Janvore, J., Mayeux, A., Fraisier, V., Nurse, P., and Paoletti, A. (2009). Spatial control of cytokinesis by Cdr2 kinase and Mid1/anillin nuclear export. Curr. Biol. 19, 961–966.

Atkins, B.D., Yoshida, S., Saito, K., Wu, C.-F., Lew, D.J., and Pellman, D. (2013). Inhibition of Cdc42 during mitotic exit is required for cytokinesis. J. Cell Biol. 202, 231–240.

Bähler, J., and Pringle, J.R. (1998). Pom1p, a fission yeast protein kinase that provides positional information for both polarized growth and cytokinesis. Genes Dev. 12, 1356–1370.

Bähler, J., Steever, A.B., Wheatley, S., Wang, Y. l, Pringle, J.R., Gould, K.L., and McCollum, D. (1998). Role of polo kinase and Mid1p in determining the site of cell division in fission yeast. J. Cell Biol. 143, 1603–1616.

Bhattacharjee, R., Mangione, M.C., Wos, M., Chen, J.-S., Snider, C.E., Roberts-Galbraith, R.H., McDonald, N.A., Presti, L.L., Martin, S.G., and Gould, K.L. (2020). DYRK kinase Pom1 drives F-BAR protein Cdc15 from the membrane to promote medial division. Mol. Biol. Cell mbcE20010026.

Bokoch, G.M. (2003). Biology of the p21-activated kinases. Annu. Rev. Biochem. 72, 743–781.

Bringmann, H., and Hyman, A.A. (2005). A cytokinesis furrow is positioned by two consecutive signals. Nature 436, 731–734.

Cabernard, C., Prehoda, K.E., and Doe, C.Q. (2010). A spindle-independent cleavage furrow positioning pathway. Nature 467, 91–94.

Celton-Morizur, S., Bordes, N., Fraisier, V., Tran, P.T., and Paoletti, A. (2004). C-terminal anchoring of mid1p to membranes stabilizes cytokinetic ring position in early mitosis in fission yeast. Mol. Cell. Biol. 24, 10621–10635.

Celton-Morizur, S., Racine, V., Sibarita, J.-B., and Paoletti, A. (2006). Pom1 kinase links division plane position to cell polarity by regulating Mid1p cortical distribution. J. Cell. Sci. 119, 4710–4718.

Chang, F., Woollard, A., and Nurse, P. (1996). Isolation and characterization of fission yeast mutants defective in the assembly and placement of the contractile actin ring. J. Cell. Sci. 109 (Pt 1), 131–142.

Chatterjee, M., and Pollard, T.D. (2019). The Functionally Important N-Terminal Half of Fission Yeast Mid1p Anillin Is Intrinsically Disordered and Undergoes Phase Separation. Biochemistry 58, 3031–3041.

Chiou, J.-G., Balasubramanian, M.K., and Lew, D.J. (2017). Cell Polarity in Yeast. Annu. Rev. Cell Dev. Biol. 33, 77–101.

Das, M., Drake, T., Wiley, D.J., Buchwald, P., Vavylonis, D., and Verde, F. (2012). Oscillatory dynamics of Cdc42 GTPase in the control of polarized growth. Science 337, 239–243.

Das, M., Wiley, D.J., Medina, S., Vincent, H.A., Larrea, M., Oriolo, A., and Verde, F. (2007). Regulation of cell diameter, For3p localization, and cell symmetry by fission yeast Rho-GAP Rga4p. Mol. Biol. Cell 18, 2090–2101.

Davies, T., Jordan, S.N., and Canman, J.C. (2016). Cell polarity is on PAR with cytokinesis. Cell Cycle 15, 1307–1308.

Endo, M., Shirouzu, M., and Yokoyama, S. (2003). The Cdc42 binding and scaffolding activities of the fission yeast adaptor protein Scd2. J. Biol. Chem. 278, 843–852.

Eng, K., Naqvi, N.I., Wong, K.C., and Balasubramanian, M.K. (1998). Rng2p, a protein required for cytokinesis in fission yeast, is a component of the actomyosin ring and the spindle pole body. Curr. Biol. 8, 611–621.

Grassetti, A.V., Hards, R., and Gerber, S.A. (2017). Offline pentafluorophenyl (PFP)-RP prefractionation as an alternative to high-pH RP for comprehensive LC-MS/MS proteomics and phosphoproteomics. Anal Bioanal Chem 409, 4615–4625.

Guzman-Vendrell, M., Baldissard, S., Almonacid, M., Mayeux, A., Paoletti, A., and Moseley, J.B. (2013). Blt1 and Mid1 provide overlapping membrane anchors to position the division plane in fission yeast. Mol. Cell. Biol. 33, 418–428.

Hayles, J., and Nurse, P. (2001). A journey into space. Nat. Rev. Mol. Cell Biol. 2, 647–656.

Hercyk, B.S., and Das, M.E. (2019). F-BAR Cdc15 Promotes Cdc42 Activation During Cytokinesis and Cell Polarization in Schizosaccharomyces pombe. Genetics 213, 1341–1356.

Jordan, S.N., Davies, T., Zhuravlev, Y., Dumont, J., Shirasu-Hiza, M., and Canman, J.C. (2016). Cortical PAR polarity proteins promote robust cytokinesis during asymmetric cell division. J. Cell Biol. 212, 39–49.

Kotak, S., and Gönczy, P. (2013). Mechanisms of spindle positioning: cortical force generators in the limelight. Curr. Opin. Cell Biol. 25, 741–748.

Kim, H., Yang, P., Catanuto, P., Verde, F., Lai, H., Du, H., Chang, F., and Marcus, S. (2003). The kelch repeat protein, Tea1, is a potential substrate target of the p21-activated kinase, Shk1, in the fission yeast, Schizosaccharomyces pombe. J. Biol. Chem. 278, 30074–30082.

Kettenbach, A.N., Deng, L., Wu, Y., Baldissard, S., Adamo, M.E., Gerber, S.A., and Moseley, J.B. (2015). Quantitative phosphoproteomics reveals pathways for coordination of cell growth and division by the conserved fission yeast kinase pom1. Mol. Cell Proteomics 14, 1275–1287.

Kettenbach, A.N., and Gerber, S.A. (2011). Rapid and reproducible single-stage phosphopeptide enrichment of complex peptide mixtures: application to general and phosphotyrosine-specific phosphoproteomics experiments. Anal. Chem. 83, 7635–7644.

Laporte, D., Coffman, V.C., Lee, I.-J., and Wu, J.-Q. (2011). Assembly and architecture of precursor nodes during fission yeast cytokinesis. J. Cell Biol. 192, 1005–1021.

Lee, M.E., Rusin, S.F., Jenkins, N., Kettenbach, A.N., and Moseley, J.B. (2018). Mechanisms Connecting the Conserved Protein Kinases Ssp1, Kin1, and Pom1 in Fission Yeast Cell Polarity and Division. Curr. Biol. 28, 84–92.e4.

Liu, J., Wang, H., McCollum, D., and Balasubramanian, M.K. (1999). Drc1p/Cps1p, a 1,3-beta-glucan synthase subunit, is essential for division septum assembly in Schizosaccharomyces pombe. Genetics 153, 1193–1203.

Loo, T.-H., and Balasubramanian, M. (2008). Schizosaccharomyces pombe Pak-related protein, Pak1p/Orb2p, phosphorylates myosin regulatory light chain to inhibit cytokinesis. J. Cell Biol. 183, 785–793.

Marcus, S., Polverino, A., Chang, E., Robbins, D., Cobb, M.H., and Wigler, M.H. (1995). Shk1, a homolog of the Saccharomyces cerevisiae Ste20 and mammalian p65PAK protein kinases, is a component of a Ras/Cdc42 signaling module in the fission yeast Schizosaccharomyces pombe. Proc. Natl. Acad. Sci. U.S.A. 92, 6180–6184.

Martín-García, R., Coll, P.M., and Pérez, P. (2014). F-BAR domain protein Rga7 collaborates with Cdc15 and Imp2 to ensure proper cytokinesis in fission yeast. J. Cell. Sci. 127, 4146–4158.

McAlister, G.C., Nusinow, D.P., Jedrychowski, M.P., Wühr, M., Huttlin, E.L., Erickson, B.K., Rad, R., Haas, W., and Gygi, S.P. (2014). MultiNotch MS3 enables accurate, sensitive, and multiplexed detection of differential expression across cancer cell line proteomes. Anal. Chem. 86, 7150–7158.

Omotade, O.F., Pollitt, S.L., and Zheng, J.Q. (2017). Actin-based growth cone motility and guidance. Mol. Cell. Neurosci. 84, 4–10.

Opalko, H.E., Nasa, I., Kettenbach, A.N., and Moseley, J.B. (2019). A mechanism for how Cdr1/Nim1 kinase promotes mitotic entry by inhibiting Wee1. Mol. Biol. Cell 30, 3015–3023.

Padte, N.N., Martin, S.G., Howard, M., and Chang, F. (2006). The cell-end factor pom1p inhibits mid1p in specification of the cell division plane in fission yeast. Curr. Biol. 16, 2480–2487.

Padmanabhan, A., Bakka, K., Sevugan, M., Naqvi, N.I., D’souza, V., Tang, X., Mishra, M., and Balasubramanian, M.K. (2011). IQGAP-related Rng2p organizes cortical nodes and ensures position of cell division in fission yeast. Curr. Biol. 21, 467–472.

Paoletti, A., and Chang, F. (2000). Analysis of mid1p, a protein required for placement of the cell division site, reveals a link between the nucleus and the cell surface in fission yeast. Mol. Biol. Cell 11, 2757–2773.

Pollard, T.D., and Wu, J.-Q. (2010). Understanding cytokinesis: lessons from fission yeast. Nat. Rev. Mol. Cell Biol. 11, 149–155.

Proctor, S.A., Minc, N., Boudaoud, A., Chang, F. (2012). Contributions of Turgor Pressure, the Contractile Ring, and Septum Assembly to Forces in Cytokinesis in Fission Yeast. Curr. Biol. 22, 1601–1608.

Ramos, M., Cortés, J.C.G., Sato, M., Rincón, S.A., Moreno, M.B., Clemente-Ramos, J.Á., Osumi, M., Pérez, P., and Ribas, J.C. (2019). Two *S. pombe* septation phases differ in ingression rate, septum structure, and response to F-actin loss. J. Cell Biol. 218, 4171–4194.

Rappaport, R., Bard, J.B.L., Barlow, P.W., and Kirk, D.L. (2009). Developmental and Cell Biology, 32. (Cambridge, GBR: Cambridge University Press) Available at: http://public.eblib.com/choice/publicfullrecord.aspx?p=4638249.

Ren, L., Willet, A.H., Roberts-Galbraith, R.H., McDonald, N.A., Feoktistova, A., Chen, J.-S., Huang, H., Guillen, R., Boone, C., Sidhu, S.S., et al. (2015). The Cdc15 and Imp2 SH3 domains cooperatively scaffold a network of proteins that redundantly ensure efficient cell division in fission yeast. Mol. Biol. Cell 26, 256–269.

Roberts-Galbraith, R.H., Ohi, M.D., Ballif, B.A., Chen, J.-S., McLeod, I., McDonald, W.H., Gygi, S.P., Yates, J.R., and Gould, K.L. (2010). Dephosphorylation of F-BAR protein Cdc15 modulates its conformation and stimulates its scaffolding activity at the cell division site. Mol. Cell 39, 86–99.

Senko, M.W., Remes, P.M., Canterbury, J.D., Mathur, R., Song, Q., Eliuk, S.M., Mullen, C., Earley, L., Hardman, M., Blethrow, J.D., et al. (2013). Novel parallelized quadrupole/linear ion trap/Orbitrap tribrid mass spectrometer improving proteome coverage and peptide identification rates. Anal. Chem. 85, 11710–11714.

Siller, K.H., and Doe, C.Q. (2009). Spindle orientation during asymmetric cell division. Nat. Cell Biol. 11, 365–374.

Snell, V., and Nurse, P. (1994). Genetic analysis of cell morphogenesis in fission yeast--a role for casein kinase II in the establishment of polarized growth. EMBO J. 13, 2066–2074.

Sohrmann, M., Fankhauser, C., Brodbeck, C., and Simanis, V. (1996). The dmf1/mid1 gene is essential for correct positioning of the division septum in fission yeast. Genes Dev. 10, 2707–2719.

Tatebe, H., Nakano, K., Maximo, R., and Shiozaki, K. (2008). Pom1 DYRK regulates localization of the Rga4 GAP to ensure bipolar activation of Cdc42 in fission yeast. Curr. Biol. 18, 322–330.

Ting, L., Rad, R., Gygi, S.P., and Haas, W. (2011). MS3 eliminates ratio distortion in isobaric multiplexed quantitative proteomics. Nat. Methods 8, 937–940.

Tojkander, S., Gateva, G., and Lappalainen, P. (2012). Actin stress fibers--assembly, dynamics and biological roles. J. Cell. Sci. 125, 1855–1864.

Vavylonis, D., Wu, J.-Q., Hao, S., O’Shaughnessy, B., and Pollard, T.D. (2008). Assembly mechanism of the contractile ring for cytokinesis by fission yeast. Science 319, 97–100.

Verde, F., Mata, J., and Nurse, P. (1995). Fission yeast cell morphogenesis: identification of new genes and analysis of their role during the cell cycle. J. Cell Biol. 131, 1529–1538.

Willet, A.H., DeWitt, A.K., Beckley, J.R., Clifford, D.M., and Gould, K.L. (2019). NDR Kinase Sid2 Drives Anillin-like Mid1 from the Membrane to Promote Cytokinesis and Medial Division Site Placement. Curr. Biol. 29, 1055–1063.e2.

Willet, A.H., McDonald, N.A., Bohnert, K.A., Baird, M.A., Allen, J.R., Davidson, M.W., and Gould, K.L. (2015). The F-BAR Cdc15 promotes contractile ring formation through the direct recruitment of the formin Cdc12. J. Cell Biol. 208, 391–399.

Wu, J.-Q., Kuhn, J.R., Kovar, D.R., and Pollard, T.D. (2003). Spatial and temporal pathway for assembly and constriction of the contractile ring in fission yeast cytokinesis. Dev. Cell 5, 723–734

Wu, J.-Q., Sirotkin, V., Kovar, D.R., Lord, M., Beltzner, C.C., Kuhn, J.R., and Pollard, T.D. (2006). Assembly of the cytokinetic contractile ring from a broad band of nodes in fission yeast. J. Cell Biol. 174, 391–402.

Yang, P., Pimental, R., Lai, H., and Marcus, S. (1999). Direct activation of the fission yeast PAK Shk1 by the novel SH3 domain protein, Skb5. J. Biol. Chem. 274, 36052–36057.

Yang, P., Qyang, Y., Bartholomeusz, G., Zhou, X., and Marcus, S. (2003). The novel Rho GTPase-activating protein family protein, Rga8, provides a potential link between Cdc42/p21-activated kinase and Rho signaling pathways in the fission yeast, Schizosaccharomyces pombe. J. Biol. Chem. 278, 48821–48830.

Ye, Y., Lee, I.-J., Runge, K.W., and Wu, J.-Q. (2012). Roles of putative Rho-GEF Gef2 in division-site positioning and contractile-ring function in fission yeast cytokinesis. Mol. Biol. Cell 23, 1181–1195.

